# Numerical Variability of functional MRI Graph Measures

**DOI:** 10.64898/2025.12.22.695524

**Authors:** Mina Alizadeh, Yohan Chatelain, Gregory Kiar, Tristan Glatard

## Abstract

Network neuroscience provides a powerful framework for studying the mechanisms underlying brain-related diseases. As analyses become increasingly computational, ensuring their numerical reliability has become a critical challenge. Small perturbations introduced during processing can propagate through complex pipelines, leading to variability in outcomes and raising concerns about the reproducibility of reported findings. Addressing this issue requires systematic evaluation of pipeline stability to ensure results remain within acceptable numerical limits. While the numerical variability of structural imaging workflows has been investigated, with findings ranging from negligible to substantial, functional MRI (fMRI) pipelines and their derived graph measures remain underexplored. Without rigorous stability assessment, conclusions drawn from these measures may remain uncertain. We systematically evaluated the numerical variability of graph measures of functional connectivity derived from the widely-used fMRIPrep pipeline and compared it to population variability. The resulting Numerical-Population Variability Ratio (NPVR) values typically ranged from 0.1 to 0.2 for most graph metrics, indicating a measurable influence of numerical variability on network-derived outcomes. NPVR values varied across brain regions, thresholding choices, and confound regression strategies. These findings highlight numerical variability as an important factor in functional network studies, particularly when examining subtle effects or working with small sample sizes.

## 1 Introduction

Variability across analytical tools and computational infrastructures can influence functional MRI (fMRI) outcomes, thereby challenging the reliability of derived findings. Large-scale studies, such as the Neuroimaging Analysis Replication and Prediction Study (NARPS) Botvinik-Nezer et al., 2020 have shown that analytical variability can substantially affect results, with teams often reaching divergent conclusions despite analyzing the same data. Analytical variability arises from the wide range of tool choices and parameter settings available for preprocessing and statistical analyses Carp, 2012; Germani et al., 2025; Li et al., 2024, as well as from differences in computational infrastructure Chatelain et al., 2024; Glatard et al., 2015.

Numerical variability, a less explored source of variability, likely plays a role in analytical variability Kiar et al., 2024. Minor computational perturbations introduced by differences in hardware architectures, operating systems, compilers, or even parallelization strategies, can propagate through analytical pipelines and amplify into meaningful discrepancies in final outcomes Glatard et al., 2015; Kiar et al., 2020; Salari et al., 2021. Such effects are particularly present in pipelines that involve optimization procedures in high-dimensional spaces, such as linear and non-linear image registration, or the training of deep-learning models.

While prior studies have examined the numerical variability of structural Mirhakimi et al., 2025 Chatelain et al., 2026 and diffusion Kiar et al., 2021 neuroimaging pipelines, its impact on functional MRI analyses remains unexplored. Given the widespread use of network neuroscience to investigate brain organization and disease, evaluating the numerical variability of network statistics is essential to ensure the validity and interpretability of findings at both the individual and group levels.

This work investigates how numerical perturbations influence the reliability of functional connectivity matrices and derived graph metrics across multiple datasets and pre-processing configurations. By quantifying the numerical variability of graph metrics, we provide insights into the robustness of network-based biomarkers and contribute to establishing best practices for ensuring computational reproducibility in neuroimaging research.

## 2 Materials and Methods

We applied the widely-used fMRIPrep preprocessing pipeline to fMRI data from the PPMI dataset to assess numerical variability, focusing on cross-sectional analyses. Numerical variability was evaluated using Monte Carlo Arithmetic (MCA) Denis et al., 2016; Parker and Langley, 1997, a technique that characterizes the sensitivity of a computational pipeline to small numerical perturbations. By repeatedly perturbing floating-point operations, MCA enables us to quantify how much variability arises purely from numerical errors. We then examined how these perturbations propagate into derived graph measures and compared the resulting numerical variability to inter-individual variability, evaluating its potential impact on downstream statistical analyses. Data was collected after approval of the local ethics 104 committees of the PPMI’s participating sites.

AI-based language tools were used solely for spelling and grammar correction. No scientific content, analysis, or interpretation was generated using AI.

### 2.1 Dataset

We used data from the *Parkinson Progression Marker Initiative* (PPMI) Marek et al., 2011, an ongoing, international, multicenter observational study designed to identify biomarkers of Parkinson’s disease (PD). From the PPMI cohort, we selected individuals with available resting-state fMRI (rs-fMRI) data, including both Parkinson’s disease (PD) patients and healthy control (HC) participants.

PPMI provides multiple rs-fMRI acquisitions using 2D gradient-echo T2*-weighted EPI with different acquisition configurations (e.g., `epi_2d`, RL, PA) and phase-encoding directions. To avoid additional variability introduced by mixing acquisition types and phase-encoding directions, we restricted our analysis to the RL runs only, including 38 of healthy controls and 147 of PD patients. The RL rs-fMRI acquisition consists of 240 volumes with 10 min total scan time, with a repetition time (TR) of 2500 ms, an echo time (TE) of 30 ms, a slice thickness of 3.5 mm, 40 axial slices, a field of view (FOV) of 224 *×* 224 mm, and a matrix size of 64 *×* 64. Participants were instructed to keep their eyes open and remain still during the scan. Since the data were in DICOM format, they were converted to NIfTI format using the HeuDiConv conversion framework (version 1.2) to enable compatibility with the preprocessing pipeline, which requires BIDS-validated input data.

### 2.2 Data Processing

All anatomical and functional data were preprocessed using *fMRIPrep* (version 23.2.1). Anatomical T1-weighted images underwent intensity non-uniformity correction, skull stripping, tissue segmentation, and nonlinear normalization to the MNI152NLin2009cAsym template.

Functional preprocessing included the generation of a BOLD reference volume, head-motion correction, and co-registration of the BOLD reference to the T1-weighted image using boundary-based registration. The corrected BOLD time series were subsequently transformed into MNI space using the composed anatomical and functional transformations. fMRIPrep also produced a comprehensive set of confound regressors, including motion parameters, framewise displacement (FD), DVARS, global signals, and components derived from aCompCor and tCompCor. All spatial resampling steps were combined into a single interpolation using the composed transforms.

Following preprocessing, functional connectomes were generated using `Nilearn`. Regional time series were extracted using a `NiftiLabelsMasker` applied to the preprocessed BOLD images, using the Schaefer 2018 parcellation with 100 cortical regions and 7 functional networks (`fetch_atlas_schaefer_2018`, *n* ROIs = 100, *n* networks = 7). Spatial smoothing (6 mm FWHM) and temporal standardization were applied during masking. To assess the influence of denoising strategies on numerical stability, two versions of the functional connectome were constructed for each subject and each MCA run:

1. **With confound regression**: time series were extracted after regressing out the six main sets of motion parameters (translations and rotations) of fMRIPrep confounds.
2. **Without confound regression**: time series were extracted without applying any confound regression, isolating numerical variability arising solely from preprocessing.

For each version, Pearson correlation matrices were computed using Nilearn’s `ConnectivityMeasure`kind=“`correlation`”), yielding one functional connectivity matrix per subject per MCA run.

### 2.3 Graph Metrics

As correlation matrices are initially complete graphs, limiting the insight from many graph metrics, they were thresholded as is commonly done, using six correlation values (0.05, 0.1, 0.2, 0.3, 0.4, 0.5) to generate binarized adjacency matrices. From these networks, we computed four local graph metrics: degree centrality, clustering coefficient, betweenness centrality, and eigenvector centrality, as well as two global metrics: small-worldness and average shortest path length. All graph measures were computed using `Python`(version 3.12) and `NetworkX`(version 3.5) library. Degree centrality quantifies the direct connectivity of a node by counting its number of adjacent neighbors, reflecting local influence. Eigenvector centrality extends this notion by assigning higher importance to nodes connected to other highly connected nodes, capturing their global prominence in the network Akbari et al., 2024. Betweenness centrality measures the fraction of shortest paths that pass through a node, indexing its role in mediating information flow. The clustering coefficient reflects functional segregation by quantifying how strongly a node’s neighbors are interconnected Prajapati and Emerson, 2021. For global topology, we computed the average shortest path length, which measures the efficiency of information integration across the entire network, and small-worldness, defined using normalized clustering and path-length relative to matched random networks, capturing the balance between segregation and integration characteristic of small-world architecture Fang et al., 2017. Together these metrics summarize a variety of information transmission qualities of the network.

### 2.4 Evaluation of Numerical Variability

Numerical variability arises from the finite-precision nature of floating-point arithmetic, where subtle rounding and cancellation errors accumulate over long computational chains. Such errors can originate from differences in elementary mathematical libraries resulting for instance from updates in operating systems, hardware, or parallelization Salari et al., 2021. Monte Carlo Arithmetic (MCA) is a technique to assess numerical stability in real-world scientific software. MCA introduces controlled, stochastic rounding noise into floating-point operations, thereby simulating the variability that would occur across diverse computational environments Parker and Langley, 1997

In this study, we used *fuzzy libmath* (FL) Salari et al., 2021, a tool that instruments the GNU mathematical library—the mathematical library used in most Linux-based systems—with MCA through the Verificarlo tool Denis et al., 2016 to inject controlled stochastic noise into elementary mathematical functions. Through dynamic linking, fuzzy libmath instruments these functions at runtime, enabling MCA perturbations without modifying or recompiling the original pipeline code.

By repeatedly executing the *fMRIPrep* pipeline using fuzzy libmath (10 MCA repetitions per subject), we obtained a distribution of perturbed outputs that captures the range of numerical variability expected across OS or library configurations. This perturbation model allows us to estimate the numerical variability inherent to the pipeline and to assess its impact on functional connectome construction and graph measures.

### 2.5 Numerical variability measures

To evaluate the impact of numerical variability on graph measures, we used the Numerical–Population Variability Ratio (*𝒱*_npv_), a framework developed by Chatelain et al., 2026. This framework quantifies numerical variability relative to variability across subjects, providing a formal comparison between the variability induced by MCA perturbations and the variability observed across subjects.

The *𝒱*_npv_ is defined as:

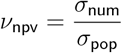

where *σ*_num_ represents numerical variability across MCA repetitions for each individual subjects and *σ*_pop_ represents inter-subject differences within each repetition. For each region of interest, *x*_*i,j*_ represents the measurement for subject *j* in MCA repetition *i*. Numerical variability quantifies intra-subject measurement consistency:

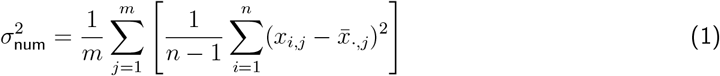

Population variability captures inter-subject differences:

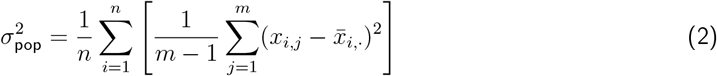

where 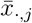 and 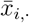 denote column and row means, *n* is the total number of MCA repetitions and *m* is the number of subjects. Higher *𝒱*_npv_ values indicate regions where computational variability potentially compromises the detection of true population differences. *𝒱*_npv_ can also be propagated to statistics used in common statistical tests. For instance, the standard deviation of Cohen’s *d* (*σ*_*d*_) can be approximated as follows Chatelain et al., 2026:

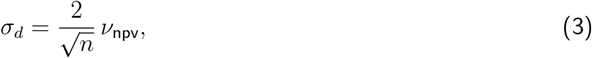

This relation formalizes how numerical variability impacts the reliability of downstream inferences. The numerical variability of other classical statistics, including the t-statistic and the F-statistic, as well as of their associated p-values, can also be expressed as a function of *𝒱*_npv_.

## 3 Results

### 3.1 The influence of numerical variability on statistical inference

To illustrate how numerical noise can influence statistical inference and experimental design, we generated a series of simulations (Figure 1). As shown in Figure 1a–b, two synthetic populations of ten subjects each were generated under “low” and “high” within-individual variability conditions with the same mean. These scenarios emulate differences in the numerical variability of derived measurements and allow us to quantify their effects on inference.

**Figure 1.**
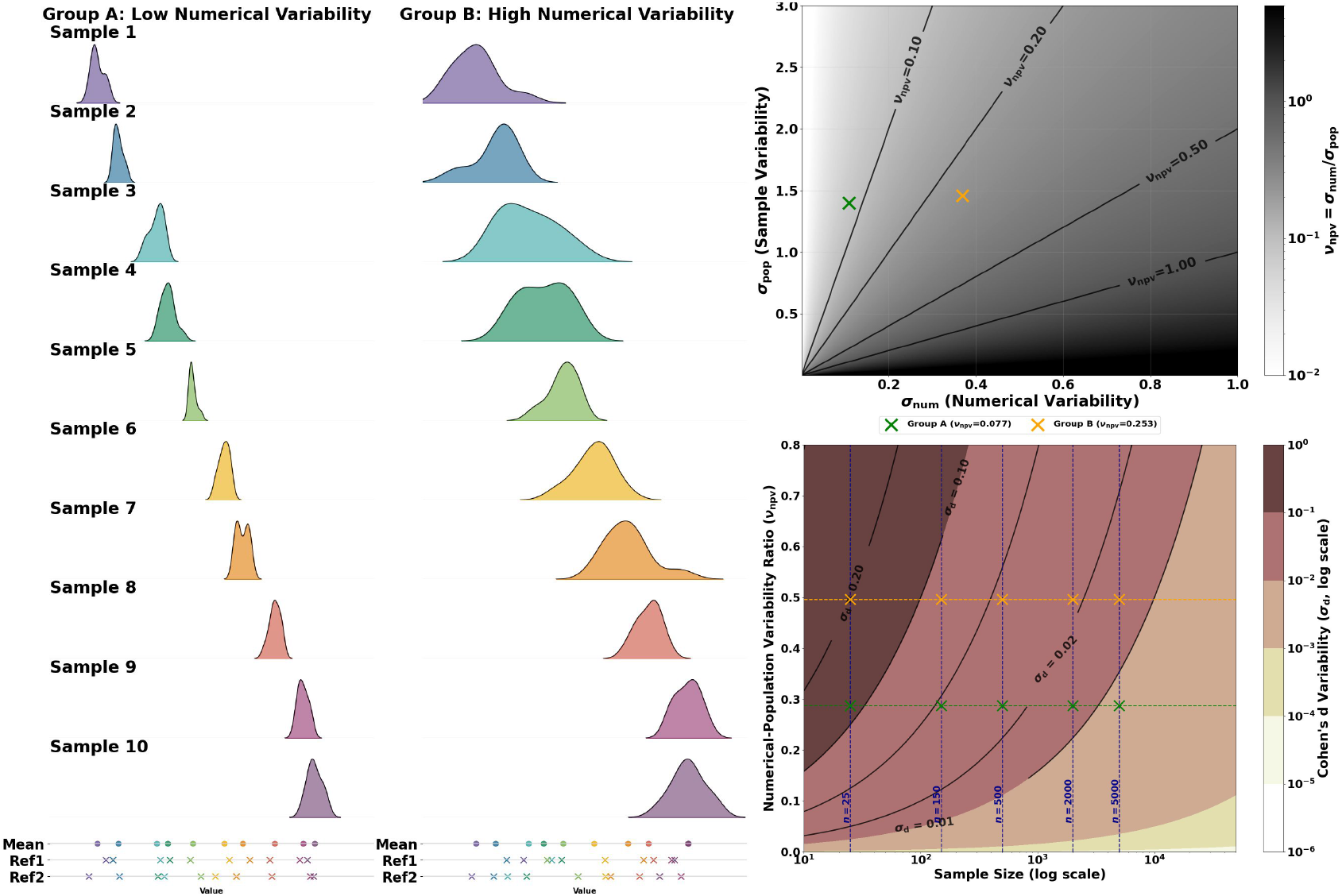
Impact of numerical variability on inference. (a–b) Simulated distributions of measurement values from two populations (10 subjects each) under low (a) and high (b) within-individual numerical variability. **(c)** Numerical-Population Variability Ratio (*𝒱*_npv_) computed for the two populations. **(d)** Effect of numerical variability on Cohen’s *d* variability (*σ*_*d*_) as a function of sample size. Markers labeled represent example sample sizes (*n* = 25, *n* = 150, and *n* = 500, respectively), included to illustrate how variability evolves with increasing *n*. Higher NPVR values (pink) correspond to greater uncertainty in the estimated effect size, which in turn requires larger sample sizes to achieve the same level of statistical reliability compared to metrics with lower NPVR (yellow).

The resulting Numerical-Population Variability Ratio (NPVR), Figure 1c captures the relative contribution of numerical variability to inter-subject variability, providing a direct measure of the impact of numerical variability

We further demonstrate how numerical variability propagates into effect size estimation (Figure 1d). Specifically, for smaller sample sizes (e.g., *n <* 100), higher NPVR values lead to substantially greater variability in Cohen’s *d* (i.e., larger *σ*_*d*_), with values ranging approximately from 0.1 to 0.5. In contrast, lower NPVR conditions yield smaller dispersion, with *σ*_*d*_ values range from 0.05 to 0.2. As sample size increases, the variability in effect size estimates diminishes for both high- and low-NPVR settings.

Together, these results make explicit the direct link between numerical variability and effect size variability. In practical terms, they show that even modest reductions in numerical variability can inflate variability in observed effects—thereby constraining both the design (e.g., required sample size) and the interpretability of neuroimaging experiments.

### 3.2 Numerical variability varies across graph statistics and network thresholding decisions

Figures 2 and 3 present the NPVR of local and global graph metrics across network thresholds. For each metric, each plot shows the mean NPVR computed across 100 nodes, along with numerical and inter-subject variability values.

**Figure 2.**
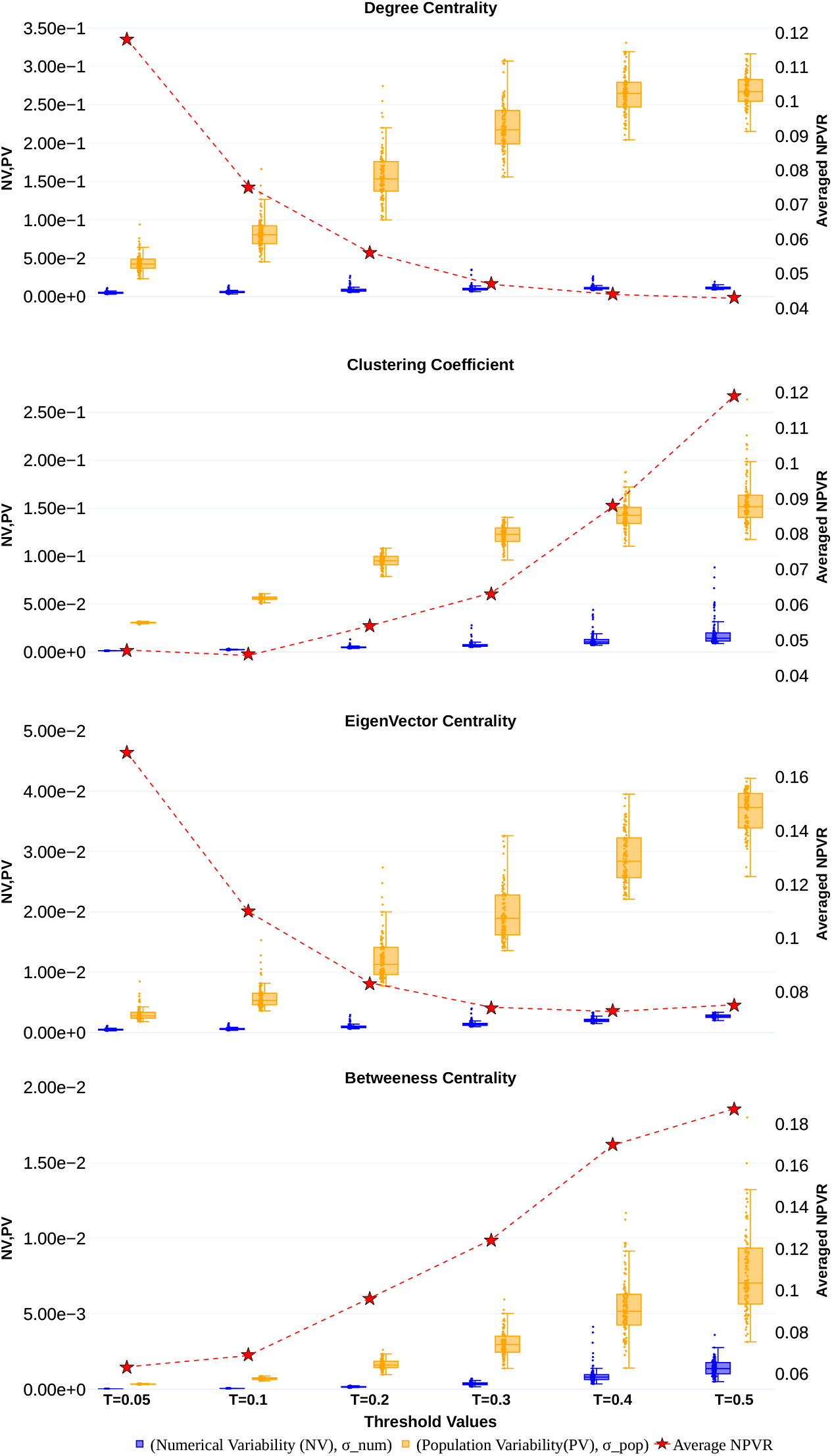
*σ*_num_, *σ*_pop_ and 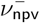 for local graph metrics across different thresholds. Each dot in a boxplot maps to the region.

**Figure 3.**
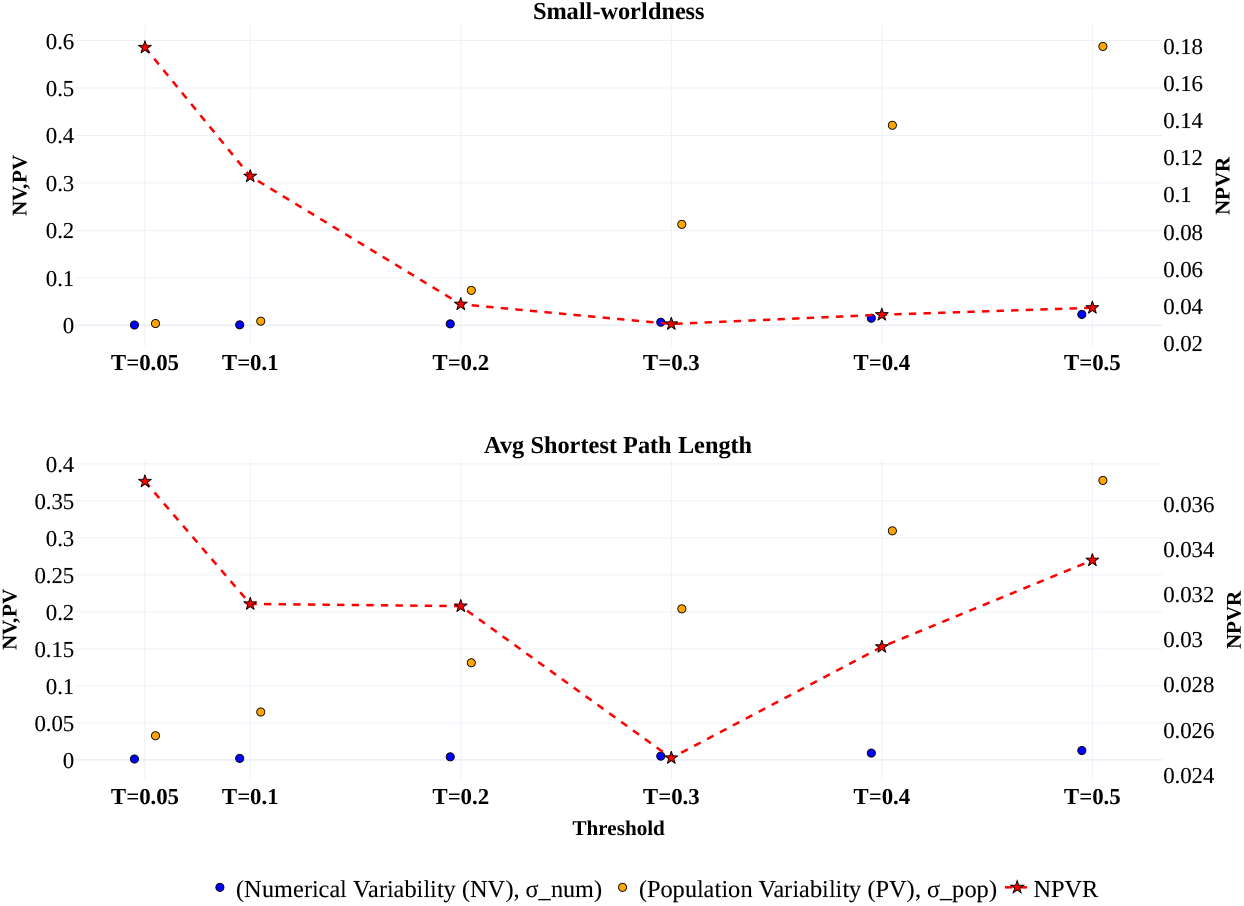
*σ*_num_, *σ*_pop_ and 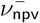 for global graph metrics across different thresholds.

#### Magnitude of NPVR

For local graph metrics (Figure 2), the average NPVR ranged from 0.04 to 0.18, while for global metrics (Figure 3), it ranged from 0.02 to 0.17, indicating that the averaged values of NPVR across regions were comparable across all metrics overall, except for average shortest path length. Among local metrics, the NPVR increased with threshold for *clustering coefficient* and *betweenness centrality*, suggesting that numerical variability grew faster than inter-subject variability. Conversely, for *degree* and *eigenvector centrality*, the NPVR decreased, indicating that inter-subject variability dominated. For global metrics, *small-worldness* and *average shortest path length* showed higher NPVR at a threshold of 0.05, followed by a decrease up to 0.3, and a subsequent rise—more pronounced for the latter.

#### Implications for downstream analyses

Although numerical variability remains smaller than intersubject variability, it may influence group-level inferences, particularly in small-sample studies. Based on Eq. 3, the observed NPVR provides a principled way to estimate how numerical variability propagates into variability in effect size estimates. For instance, a NPVR of 0.18 for *betweenness centrality* at threshold 0.5 implies that achieving stable estimates of Cohen’s *d* (variance of Cohen’s d range (10^*−*3^, 10^*−*2^)) may require sample sizes exceeding 1000 subjects. Thus, even modest numerical perturbations may interact with the small effect sizes and limited sample sizes typical of network-neuroscience studies Koen et al., 2023, potentially influencing the detectability and replicability of reported findings. These results emphasize the need to account for numerical variability when interpreting group differences, especially in analyses targeting weak effects.

### 3.3 Regional variation in NPVR interacts with threshold decisions

Figure 4 shows regional maps of the NPVR (numerical-to-inter-subject variability) for the four local graph metrics across thresholds. Overall, the NPVR is spatially heterogeneous and depends both on the metric and on the applied threshold. For lower thresholds, *degree centrality* and *eigenvector centrality* exhibit slightly higher NPVR values with increased variability across regions. In contrast, for *clustering coefficient* and *betweenness centrality*, higher thresholds tend to produce substantially larger NPVR magnitudes, accompanied by pronounced regional outliers. The spatial variability is also greater for these metrics at higher thresholds, reflecting more localized and extreme numerical variability.

**Figure 4.**
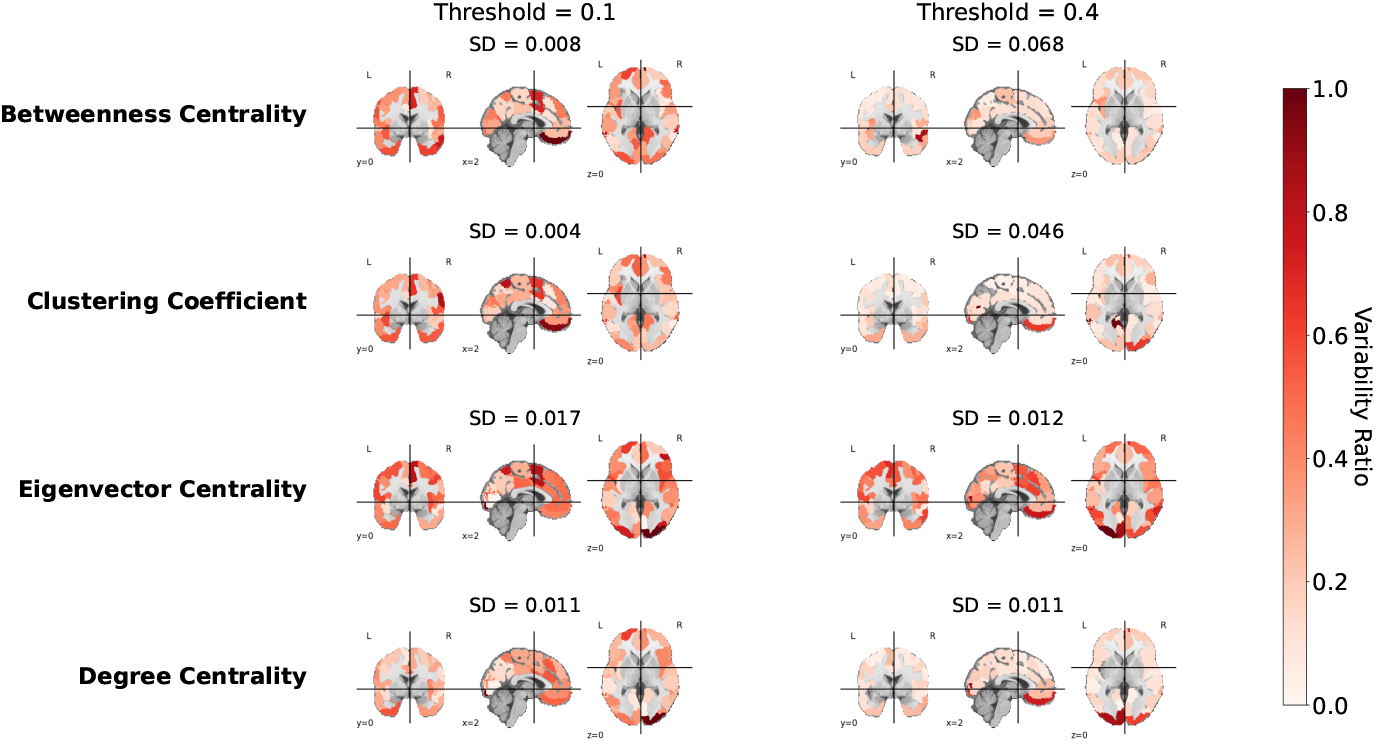
Normalized Regional NPVR maps across metrics and thresholds. Warmer colors indicate higher normalized NPVR, reflecting greater relative contribution of numerical variability to total variability.

This result highlights that numerical variability is not uniformly distributed across brain regions and that certain regions or metrics are more sensitive to thresholding. Consequently, inference drawn from regional graph metrics—such as node-level comparisons or correlations with behavior—may be differentially affected by numerical variability depending on both the metric and the chosen threshold.

### 3.4 Trade-offs of Using Confound Removal

To assess the influence of confound regression on the numerical variability of the graph metrics, we compared the results obtained from the *with-confound* and *no-confound* pipelines. In all panels of Figures 5 and 6, the plotted values represent the difference between the two processing pipelines, calculated as *no-confound* minus *with-confound*. These differences are shown across thresholds for numerical variability, population variability, and NPVR, the average ratio across regions and the global graph metrics.

**Figure 5.**
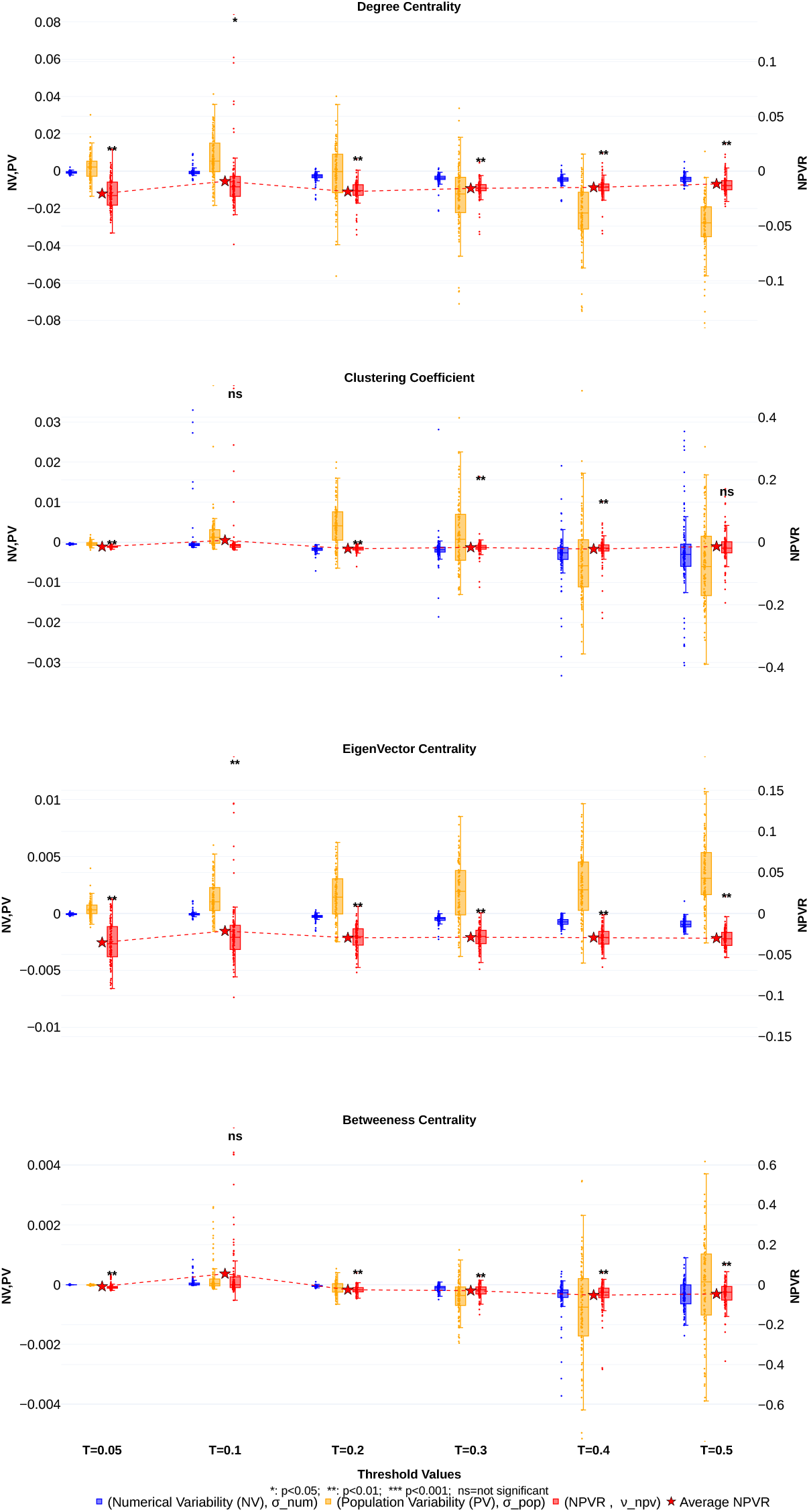
Differences of Population, Numerical variability, and NPVR between the *no-confound* and *with-confound* for Local Graph Metrics

**Figure 6.**
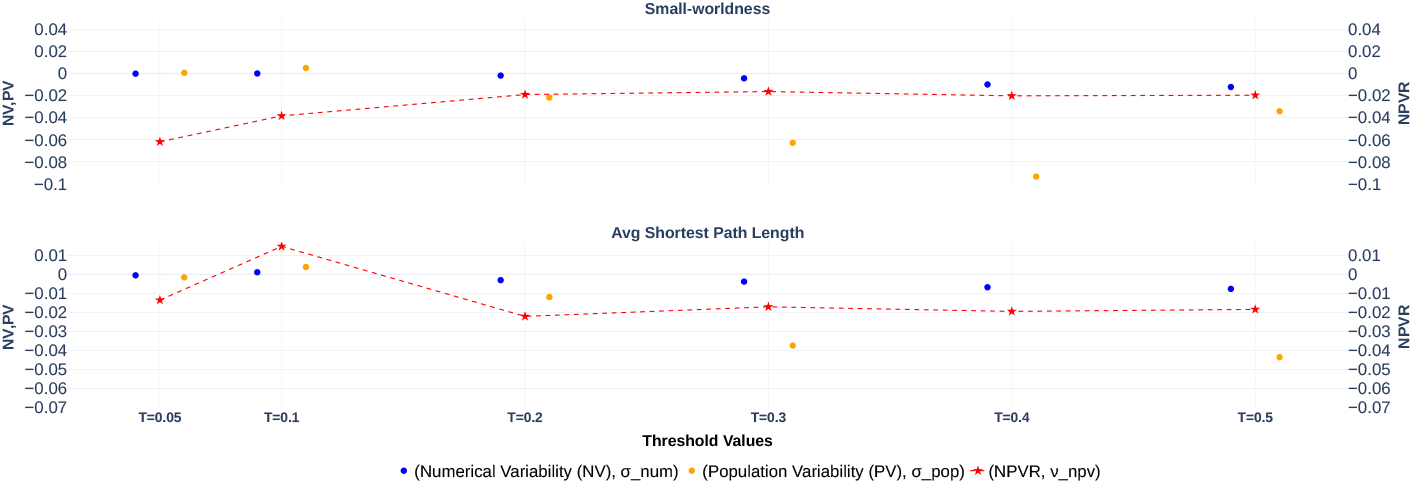
Differences of Population, Numerical variability, and NPVR between the *no-confound* and *with-confound* for Global Graph Metrics

As seen in Figure 5, the no-confound pipeline generally preserved the same overall trend in average NPVR across thresholds as the with-confound pipeline. The difference plots of the NPVR (purple boxplots) are predominantly negative, as are the differences in region-averaged NPVR (red stars), suggesting lower numerical variability in the *no-confound* pipeline relative to the *with-confound* pipeline across most metrics and thresholds. Permutation tests applied to each NPVR distribution confirmed that these differences were statistically significant and consistently negative, supporting the conclusion that confound regression increases numerical variability in this setting. However, this reduction in numerical variability does not necessarily imply better validity: removing confound regression may complicate the interpretation of the resulting connectivity estimates and could compromise the accuracy of the derived functional connectomes.

## 4 Discussion

The reliability of quantitative network neuroscience depends not only on accurate mapping of functional connectivity but also on the stability of the derived measures. While methodological discussions typically emphasize sample size, biological variability, and effect size, measurement noise from computational sources remains largely unaddressed. We showed that numerical variability originating from floating-point arithmetic can influence functional connectomes and graph metrics. Through the Numerical-Population Variability Ratio (NPVR), we quantified this computational noise relative to inter-subject differences, framing numerical variability in statistical terms commonly used to evaluate biological effects.

Our findings reveal that numerical variability is not uniform and varies across metrics, thresholds, and spatial locations. However, the observed average NPVR values consistently range between 0.1 and 0.2, indicating that numerical variability accounts for a measurable fraction of the variability observed within a population. This suggests that numerical variability may interfere with downstream analyses, particularly in studies with small sample sizes and when measuring small effect sizes, making its impact more substantial. In typical case–control connectivity studies, group sizes range from approximately 15 to 73 individuals (10th to 90th percentiles; median sample size *∼* 30 subjects per group), consistent with broader trends in neuroimaging Koen et al., 2023. Even modest numerical variability on the order of *𝒱* _npv_ = 0.1 can introduce variability in Cohen’s *d* effect size estimation in the range of [0.01, 0.1], as shown in Figure 1c. When examining subtle effects around *d*_true_ *≈*0.10, a variability of *σ*_*d*_ = 0.05 results in variability amounting to 50%of the target effect size, potentially masking or distorting true group-level differences.

We observed similar NPVR value ranges in structural MRI Chatelain et al., 2026, suggesting that this interval may be a good ballpark estimate of numerical variability in MRI analyses. This range provides a useful reference point for contextualizing results in the current literature and for approximating the variability that propagates into downstream analyses, including its impact on statistical inferences such as effect sizes, F-statistics, and other measures as proposed by Chatelain et al., 2026.

However, the specific NPVR values reported in our study will likely vary across multiple factors. Numerical variability is inherently shaped by the chosen processing pipeline, the task or resting-state paradigm, sample characteristics, and the neurobiological properties of the studied population. Prior work has already demonstrated that certain preprocessing steps introduce substantial numerical instability: for example, Mirhakimi et al., 2025 showed that linear registration exhibits marked numerical variability across runs, and Gonzalez-Pepe et al., 2025 demonstrated that the FastSurfer deep-learning pipeline does not yield lower numerical variability than the traditional FreeSurfer workflow. These findings reinforce that numerical variability is not tied to a specific algorithmic class and may emerge across both conventional and machine-learning–based pipelines. Understanding how NPVR interacts with specific preprocessing steps, graph-construction choices, or distinct subject cohorts may ultimately become informative in its own right. In particular, if certain pipelines or populations consistently exhibit elevated NPVR in specific regions or metrics, this could motivate novel quality-control strategies or even the development of computational biomarkers. A systematic investigation of these interactions would therefore be an important direction for future work.

Moreover, we focused on cross-sectional studies, where numerical variability is evaluated relative to intersubject differences. How longitudinal pipelines behave in fMRI analysis remains an open question and an important direction for future research.

Approximately 80%of network studies apply some form of thresholding to the connectivity matrix, making it common in both structural and functional network analyses. Finding the optimal thresholding approach remains an open question Koen et al., 2023.

Future work could explore how network sparsification and different thresholding methods influence the NPVR of graph measures. Our results suggest that an intermediate threshold (around 0.2) consistently minimizes NPVR across metrics.

Other post-processing choices also influence the distribution of numerical variability. Omitting confound regression consistently lowers NPVR, but this does not necessarily imply more accurate estimates. Numerical stability is only one dimension of pipeline performance: a method can be highly stable yet produce inaccurate results.

Reported NPVR means can be leveraged to evaluate how processing choices interact with expected effect sizes and available sample sizes, thereby informing processing decisions and the selection of thresholds that are more likely to yield robust findings

Similarly, factors such as atlas selection should be considered from the perspective of numerical variability. As noted in Koen et al., 2023, more detailed atlases introduce more parallel tests and may generally show greater measurement error in observed connectivity.

Overall, our results position numerical variability as an important factor in network studies, especially when investigating subtle effects or working with small sample sizes.

## 5 Conclusion

In this study, we examined the numerical variability of fMRI-derived graph metrics using data from the Parkinson’s Progression Markers Initiative (PPMI) cohort. We quantified numerical variability relative to inter-subject differences using the Numerical–Population Variability Ratio (NPVR). Across graph metrics, NPVR values typically ranged between 0.1 and 0.2, indicating that numerical variability accounts for a non-negligible fraction of population-level variability. NPVR can be propagated to statistics commonly used in group-level analyses, such as the variability of effect size estimates. In particular, higher NPVR values combined with the small sample sizes typically used in neuroimaging studies can substantially increase variability in effect size estimates, thereby compromising the detection and interpretation of true population-level effects. These findings underscore the need to explicitly account for numerical variability as an important source of variability affecting both derived graph measures and subsequent statistical inferences.

## Data and Code Availability

The unprocessed dataset is available through the Parkinson’s Progression Markers Initiative (PPMI) website (https://www.ppmi-info.org/access-data-specimens/download-data). Due to data-use restrictions imposed by PPMI, the raw and derived data cannot be publicly shared; however, they may be obtained upon submission of a request and compliance with the PPMI Data Usage Agreement.

DICOM images were converted to NIfTI format, as required for fMRIPrep inputs, using *HeuDiConv* (version 1.2) on the *Narval* cluster hosted by *Calcul Québec*, which is part of the *Digital Research Alliance of Canada*. The Python script and notebooks used for this study are available in the GitHub repository: https://github.com/mina94az/Numerical-Variability-of-functional-MRI-Graph-Measures.

Floating-point perturbations were introduced using the *fuzzy-libmath* instrumentation provided by the Verificarlo framework. The corresponding documentation and container image are publicly available at: https://github.com/verificarlo/fuzzy/tree/master#fuzzy-libmath.

All preprocessing was performed with fMRIPrep (version 23.2.1) instrumented with *fuzzy-libmath*, on the *Narval* cluster hosted by *Calcul Québec*.

Functional connectomes were generated using Python scripts available in the GitHub repository and were executed on the *Rorqual* cluster hosted by *Calcul Québec*, using Python 3.12 and the NetworkX (version 3.5) library. Downstream analyses were performed locally on a computer running Ubuntu 22.04.5 LTS, using Python 3.11.

## Author Contributions

**Conceptualization:** MA, YC, GK, TG; **Data curation:** MA; **Formal analysis:** MA, YC, GK, TG; **Funding acquisition:** TG; **Investigation:** MA; **Methodology:** MA, YC, GK, TG; **Project administration:** MA, GK, TG; **Supervision:** GK, TG; **Validation:** MA, YC, GK, TG; **Visualization:** MA; **Writing – original draft:** MA; **Writing – review & editing:** MA, YC, GK, TG.

## Declaration of Competing Interests

The authors have declared that no competing interests exist.

## Acknowledgments

M.A. was supported by a bursary funded through an NSERC Discovery Grant awarded to T.G., entitled *“Numerical Stability in Data Science”*.

## Notes

### Competing Interest Statement

The authors have declared no competing interest.

### Summary of Updates

updating figures, acknowledgment, and adding a conclusion

